# A topographical and physiological exploration of C-tactile afferents and their response to menthol and histamine

**DOI:** 10.1101/2021.07.02.450784

**Authors:** Line S. Löken, Helena Backlund Wasling, Håkan Olausson, Francis McGlone, Johan Wessberg

## Abstract

Numerous microneurography studies in the human peroneal nerve have suggested that CT afferents are lacking in the more distal parts of the limbs. Here we recorded from unmyelinated low-threshold mechanosensitive afferents in the peroneal and radial nerves, with the most distal receptive fields located on the proximal phalanx of the third finger for the superficial branch of the radial nerve, and near the lateral malleolus for the peroneal nerve. We found that the physiological properties with regard to conduction velocity and mechanical threshold, as well as their tuning to brush velocity, were similar in CT units across the antebrachial, radial and peroneal nerves. Moreover, we found that while CT afferents are readily found during microneurography of the arm nerves, they appear to be much more sparse in the lower leg compared to C nociceptors. We continued to explore CT afferents with regard to their chemical sensitivity and found that they could not be activated by topical application to their receptive field of either the cooling agent menthol or the pruritogen histamine. In light of previous studies showing the combined effects that temperature and mechanical stimuli have on these neurons, including a lack of responsiveness to heat, these findings add to the growing body of research suggesting that CT afferents constitute a unique class of sensory afferents with highly specialized mechanisms for transducing gentle touch.

## INTRODUCTION

Since the discovery that the skin of man is innervated with unmyelinated tactile afferents (CT) that convey social and emotional aspects of touch (Löken et al. 2009; Olausson et al. 2002; Vallbo et al. 1999; Wessberg et al. 2003), extensive explorations of the physiology and function of these afferents have been made (McGlone et al. 2007; McGlone et al. 2014; Morrison et al. 2010; Olausson et al. 2010). Despite this, aspects of the properties of CT afferents as well as their prevalence and density in different areas of the human body have remained elusive. The earliest observations of the presence of CTs in human skin were made by Johansson et al. (Johansson et al. 1988) during microneurography recordings in the supra- and infraorbital nerves and two years later, Nordin described the physiological properties of CTs in these nerves in much finer detail (Nordin 1990). The finding that CT afferents showed a vigorous response to stroking with a soft cotton swab was particularly intriguing, as C-fibers were generally thought to convey temperature, itch and nociception. Because such a large body of microneurography studies had been performed on the nerves of the arms and legs (Hagbarth et al. 1972; Jorum et al. 1989; Vallbo and Johansson 1984), the data presented by Nordin was initially thought to be specific for the face. This turned out not be true. Shortly thereafter, during microneurographic registrations from the lateral and dorsal antebrachial cutaneous nerves (Vallbo et al. 1993; Vallbo et al. 1999), CT afferents were found to be densely represented in the nerves of the arm as well. In addition, CT afferents were later also observed in the lateral cutaneous femoral nerve, innervating the thigh (Edin 2001).

The physiological characterizations of CT afferents have shown that they display characteristics such as fatigue, afterdischarge activity, intermediate adaptation, and sensitivity to mechanical stimuli and cooling stimuli (Ackerley et al. 2018; Loken et al. 2009; Nordin 1990; Vallbo et al. 1993; Vallbo et al. 1999; Wessberg et al. 2003). These properties, and in particular their poor temporal coding abilities and high activity during slow stroking movements, led to the proposal that CT afferents constitute a separate pathway mediating affective touch in the body (Olausson et al. 2002; Vallbo et al. 1999). This proposal was solidified by the finding that there is a negative quadratic relationship between brush velocity and mean firing rate, which is mirrored in perceptual ratings of pleasantness in response to the same stimuli (Loken et al. 2009) and other smooth materials (Essick et al. 1999). Furthermore, the tuning to slow touch that CT afferents display is strongest at temperatures that mimic skin-to-skin contact (Ackerley et al. 2014). CT afferents are clearly separable from C mechano-nociceptors by having mechanical threshold of 2.5 mN or lower (Vallbo et al. 1999). Furthermore, CT afferents are insensitive to heat (Nordin 1990; Vallbo et al. 1999) which separates them from the large class of C fibers that are peptidergic (Basbaum et al. 2009). Analysis of the responsiveness to cooling and combined mechanical and temperature stimuli have revealed that they do not appear to signal cooling alone, but that the combination of cooling and mechanical stimulation may alter their firing properties (Ackerley et al. 2018; Nordin 1990).

Despite these extensive investigations, fundamental questions remain regarding the distribution of CT afferents in the more distal parts of the limbs, and their sensitivity to topically or transdermally delivered chemical compounds. Recent studies have shed further light on the gene expression that may be underlying specific transduction mechanisms in mouse CLTMRs (the mouse equivalent of CTs)(Usoskin et al. 2015). It is therefore of increasing interest to establish the degree to which rodent CLTMRs and human CTs have shared characteristics as this opens up possibilities for translational studies. Studies in mice also describe CLTMs as innervating specific types of hair follicles and although this may be similar in human skin it is also known that the epidermis is densely innervated by free nerve endings, some of which may well be CTs, and therefore an easy target for topical ligands (Kennedy and Wendelschafer-Crabb 1993). We therefore set out to investigate the presence of CT afferents in different cutaneous nerves as well as how their respective physiological properties compare across skin sites. Lastly, we investigated the responsiveness of CT afferents to histamine, a well-known pruritogen (Simone et al. 1991), and menthol, a ligand for the TRPM8 cold-sensitive receptor (Bautista et al. 2007; de la Pena et al. 2005; Dhaka et al. 2006; Dhaka et al. 2007).

## METHODS

### Participants

The participants were recruited through advertisements posted in the university and mainly consisted of students in the medical department. Forty-five participants were included in the study that were required to be healthy, with no neurological illness. The mean age was 24 years (range 20-31) and 13 were male. Written, informed consent was acquired before commencing experiments and a financial compensation was given for their time. The University of Gothenburg ethics committee approved the experimental protocol that was performed in accordance to the declaration of Helsinki.

### Experimental procedure

Using the microneurography technique (Vallbo and Hagbarth 1968) we recorded from single CT afferents in the antebrachial and radial nerves that innervate the arm, and from CT and C nociceptors in the peroneal nerve which innervate the leg. Participants were seated comfortably in an adjustable dental chair with their left arm or leg supported by a vacuum airbag for stability. A custom-made pre-amplifier (Department of Physiology, Umeå University) and a silver-plated ground plate was attached to the participants’s forearm or leg.

### Search procedure

We used an electrical search procedure to locate the nerves. For the peroneal nerve the electrode was inserted below the fibular head, for the antebrachial nerve at the level of the elbow and for the radial nerve at the dorsal aspect of the forearm. The skin was palpated to find the ideal place for insertion of the stimulating and recording electrodes, and an uninsulated reference electrode was inserted ∼5 cm from this site. The stimulating electrode was uninsulated (35 or 50mm length, 200 μm shaft diameter, ∼5 μm tip diameter; FHC, Bowdoin, ME) and was used to deliver 200 ms-square, negative, 1-Hz pulses at low current until the participant reported paresthesia in the innervation area. Once an ideal electrode position was obtained, the depth and angle of the electrode was noted and withdrawn slightly from its proximity to the nerve. A new electrode was then inserted at the noted depth and angle distal to the search electrode. Recordings were made from single afferents. When the tip had attained an intrafascicular position, the experimenter searched for single units by lightly stroking with his/her fingertips over the skin on the surface of the innervation area and making minimal adjustments of electrode position. Single units that were identified as unmyelinated (by negative deflection and latency), responded to soft brush stroking and had amplitudes distinct from the noise were further studied. The nerve signal was recorded at 12.8 kHz with a passive band-pass filter set to 0.2–4.0 kHz and stored on a PC using the ZOOM/SC system developed at the Department of Physiology, Umeå University, Sweden. Recorded nerve impulses (spikes) were inspected offline on an expanded time scale using in-house software implemented in MATLAB (The Mathworks, Natick, MA) and were accepted for subsequent analyses only if they could be validated as originating from a single afferent.

### Unit identification

We identified CT afferents according to the criteria set in previous studies (Vallbo et al. 1993; Vallbo et al. 1999; Wessberg et al. 2003). First, conduction velocity was estimated from the response latency to mechanical tap stimulation using a hand-held strain gauge device. Distinct taps with a blunt probe were delivered toward the most sensitive spot within the receptive field and the minimal latency from indentation to unit response was used to estimate conduction velocity. Myelinated afferents (conduction velocity > 2 m/s) were also studied but are not further described here. Thresholds to mechanical stimuli were assessed using von Frey monofilaments, and defined as the weakest stimulation force that the unit consistently responded to. Unmyelinated afferents (conduction velocity < 2 m/s) were classified as CT afferents if they displayed a clear response (several impulses) to low mechanical threshold monofilament bristles below 2.5mN, and vigorous response to brush stroking. Unmyelinated afferents were classified as nociceptors if they displayed a high mechanical threshold to monofilament bristles (>5mN) and no response to brush stroking. For all CT afferents, conduction velocity was calculated based on response to distinct taps with a strain gauge. Conduction velocity in nociceptors was measured in the same fashion as well as with electrical stimulation, which confirmed the accuracy of the strain gauge measures.

### Experiment

#### Mechanical stimuli

Units were initially explored with a number of handheld mechanical stimuli such as gentle touch by finger stroking across the receptive field, wooden sticks and a hand-held soft watercolor brush stroked across the receptive field. The presence of after-discharge, which may occur upon initial stimulation, was noted in the protocol. For some units we also tested the response to long-lasting indentation with a suprathreshold von Frey filament (see results for number of units tested in this way). The timing of these hand-held stimuli was indicated with a foot pedal.

### After identification of a CT unit, we applied one or more of the following tests

#### Robotic Tactile Stimulator (RTS)

Stimulation was made as in (Loken et al. 2009). In short, we used an artist’s flat, soft watercolor brush made of fine, smooth, goat’s hair (Vang size 18, type 43718, Oskar Vangerow, Ottobrunn, Germany). The bristles were 22 mm long, and the width of the brush was 20 mm. Following unit identification brush stroking was applied by means of a custom built robotic tactile stimulator (RTS) (Dancer Design, Saint Helens, UK) that produced brush stroking with velocities of 0.1, 0.3, 1, 3, 10 or 30 cm/s. The brush was moved over the skin in a rotary fashion by a DC motor (Maxon Motor AG, Sachseln, Switzerland) fitted with a reduction drive and position encoder. A 6-axis force/torque transducer (ATI Industrial Automation, Apex, NC) was mounted between the shaft of the DC motor assembly and the hub, which held a probe and brush. The DC motor and transducer assembly was mounted on a linear drive, driven by a step motor (Parker Hannifin Corp., Rohnert Park, CA). Both the DC and step motors were under computer control. Stimuli of different velocities were applied in randomized order with an inter-stimulus pause of 30 s for CT afferents (to avoid fatigue). In experiments where histamine or menthol was applied, we used 10 second inter-stimulus intervals when testing responsiveness to brush due to the time constraints inherent in these long recording protocols.

#### Menthol

After localization and characterization of CT units as described above, recording commenced and the receptive field was first stimulated with a hand-held brush. Subsequently, we recorded the normal response to brushing using the RTS. The direction of brush strokes was set from proximal to distal and with normal force at calibration 0.2 N. Slow and fast brush strokes (1 and 10 cm/s) were delivered at intervals of 10 seconds. This protocol was repeated four times followed by recording and application of ethanol for 5 minutes. Subsequently, the brush stimulation was repeated as above after which a pad of menthol solution (30% in ethanol) was applied on the skin for 5 minutes. We asked the participants to report if there was any sensation of cooling and then repeated the mechanical stimulation protocol again. The recording continued while asking the participant about the sensation of cooling at intervals of 60 seconds during 5 minutes. The skin was then cleaned with ethanol and water. Nerve recordings were maintained throughout the experiment until well after the skin had been cleaned.

#### Histamine

After localization and characterization of CT units as above, recording commenced and the receptive field was stimulated with a hand-held brush. We alternated application of fast and slow strokes (1 and 10 cm/s) with 10 seconds intervals. The incidence of brush strokes were noted in the recording and the procedure was repeated 3 times. We then recorded baseline with no stimulus for 2 minutes. Subsequently a drop of saline was applied to the receptive field and following this we applied Iontophoresis, 100 µA (= 1 mA, 10%) for 10 seconds. After 2 minutes, the saline was wiped off and brush strokes were again applied as above. After these baseline measures, we applied a drop of histamine (1%) and iontophoresis commenced, 100 µA (= 1 mA, 10%) for 10 seconds. We waited 2 minutes and noted the development of flare and wheal. If flare and wheal were missing, the iontophoresis procedure was repeated, otherwise the receptive field was wiped with saline and fast and slow brushing was repeated three times followed by a 2-minute recording without stimulation. We noted the development of itch, flare and wheal, the time of their peak, and subsequent attenuation throughout this procedure. In conjunction with noting the development and attenuation of the flare and wheal, participants were asked to report the qualitative sensation and specifically if there was any sensation of itch. We documented the participant’s verbal reports starting at 2 minutes post histamine iontophoresis and continuing at 3-minute intervals for 20 minutes following the histamine iontophoresis.

### Analysis

The mean firing rate was calculated from the mean of the shortest inter-spike intervals. Firing rates were then reported as mean and standard error of mean (SE) for each individual unit. Parametric tests were not used due to low sample size using with the full protocol. Regression analysis for curve fit was done by transforming velocity, the independent variable, to log10 values. Calculations were done in MATLAB and SPSS.

## RESULTS

CT afferents (lateral antebrachial nerve: n=27/32 experiments, radialis n=8/9 experiments, peroneus n=4/17 experiments) were identified by a low mechanical threshold to monofilament bristles (lateral antebrachial nerve n=27, mean threshold 0.85mN, range 0.04-2.5mN; radial nerve n=8, median threshold 0.68 mN, range 0.27-2.5mN; peroneal nerve n=4, median threshold 1.6 mN, range 0.7 – 1.6 mN), slow conduction velocity (lateral antebrachial median 0.9m/s, radialis median 0,98 m/s, peroneus median 1 m/s, range 0.9-1.1m/s m/s), and vigorous response to brush stroking. The majority of units in the sample recorded from the lateral antebrachial nerve have been described previously with respect to their response to brush velocity (Loken et al. 2009) and one unit from the radial nerve was included in a previous publication (Watkins et al. 2021). The most distal receptive fields were located on the proximal phalanx of the third finger for the superficial branch of the radial nerve, and ∼6 cm proximal to the lateral malleolus for the peroneal nerve. C nociceptors were identified by high threshold >5mN indentation with monofilaments and none of the units responded to a soft brush stroke. One unit responded with a few impulses to finger stroking of the receptive field (n=18, threshold range 5.4-59mN, mean 27mN, mean conduction velocity 0.9 m/s).

### Distribution of CT units across the skin

We performed an extensive search for CT units in the peroneal nerve and obtained an intrafascicular position for stable recording in 17 of these experiments. In the peroneal nerve, background sympathetic activity was commonly recorded, making identification of C units more challenging, which in contrast is unusual when recording from the radial or lateral antebrachial nerves. We used a similar search technique as for the nerves of the arm (i.e. stroking the skin while slowly adjusting the recording electrode), and in addition we applied pinching to the skin to identify high threshold C fibers. In this sample we found 5 times more C high-threshold fibers (n=20) compared to C low-threshold afferents (n=4). In relative terms the detection of a CT unit in the peroneal nerve was much less common than for the nerves of the arm (4 units from 17 peroneal experiments versus 35 units from 41 experiments for the forearm nerves). We marked the location of all recorded CT units across the arms and legs for the lateral antebrachial, radial and peroneal nerve as well C-nociceptors (see Figure 1 A-D). In order to explore the response to brush stroking in the different nerves we used the RTS. The setup for brush stroking across the receptive field of units in the lower leg is shown in Figure 1E.

**Figure 1.**
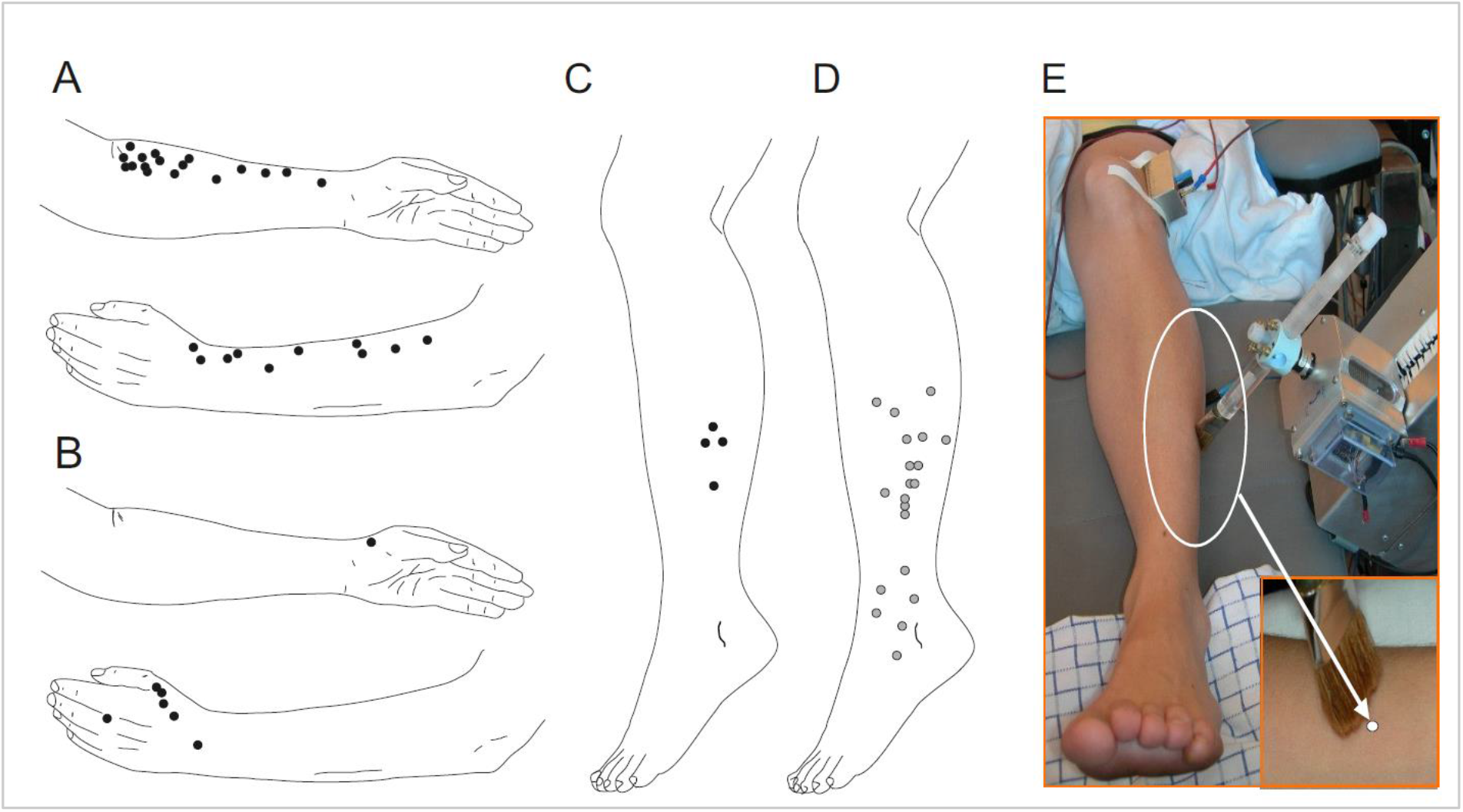
Distribution of CT units across the skin. Dots mark the location of all recorded CT units on arms (A, B) and legs (C) and nociceptors marked in (D). The setup and brush stimulus for the peroneal nerve recordings is shown in (E). The recordings for the lateral antebrachial nerve consisted of 27 experiments, the radial nerve 9 experiments and the peroneal nerve 17 experiments.

### Mechanical response properties in units across nerves

We compared basic properties of C-tactile afferents across the different nerves. We have previously shown the response to brush stroking at velocities of 0.1-30 cm/s in the lateral antebrachial nerve (Loken et al. 2009). Here we explored two of the four peroneal nerve units in the same fashion and they showed similar response properties to brushing as described for CT units of the arm (Fig 2A-B shows single stroke data from a peroneal nerve unit). A fast brush stroke (30 cm/s, Fig 2A) typically only evoke a couple of impulses in these units while a slow brush stroke (3 cm/s, Fig 2B) typically evokes a vigorous response. None of the four CT units in the peroneal nerve exhibited afterdischarge during our recordings. Four out of 20 nociceptors recorded from the peroneal nerve displayed after-discharge in response to a mechanical tap by the strain gauge upon initial exploration (supplementary figure 1). Afterdischarge is a relatively common feature in CT units of the arm and appeared in 11 out of 35 units. A clear example is shown in figure 2C where the unit, recorded from the lateral antebrachial cutaneous nerve, repeatedly fired with an extensive tail of impulses for several seconds after the stimulus had left contact with the skin. The subject could not report of any particular sensation in conjunction with this phenomenon, and its functional relevance remains unknown. We also tested delayed acceleration in all CT units of the peroneal nerve. One afferent, recorded from the peroneal nerve, showed a delayed acceleration of impulse response to sustained monofilament indentation (45 mN) (figure 2D). The initial few seconds of adaptation was followed by a period of low activity for 20 s, after which firing increased markedly again for about 90 s. This phenomenon, called delayed acceleration, has previously been reported in some CT units in the lateral antebrachial cutaneous nerve (Vallbo et al. 1999). In the peroneal nerve, out of 3 units tested with sustained indentation, 1 unit showed a delayed acceleration response (figure 2 D). In the arm experiments we found 2 units out of 7 tested that showed delayed acceleration. Note that the timescale is long in this figure and that individual spikes cover the view almost completely of the background. Individual spikes are superimposed in inset. The tuned response to intermediate brush velocity shown for CT afferents previously was consistent across units in the different nerves. Figure 2 E-F shows single unit examples in response to the full brush stroking protocol (0.1-30 cm/s) in the lateral antebrachial (E), radial (F) and peroneal (G) nerve. Although some variability is present in individual units as a response to the brush, the inverted u-shape curve (quadratic fit) of firing rate as a function of velocity is highly consistent.

**Figure 2.**
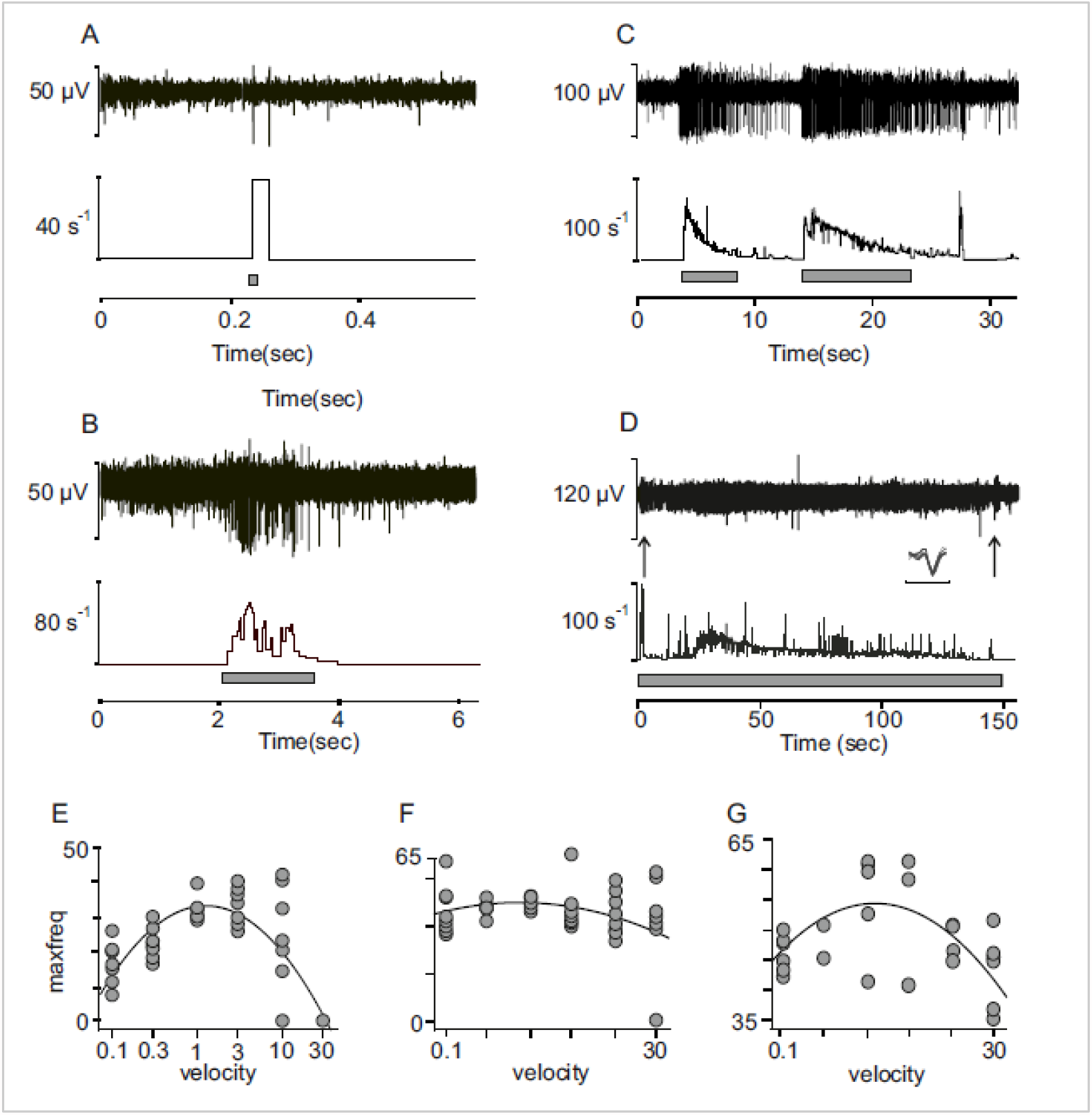
Mechanical (brush) response properties in units across nerves. A-B) Spikes in response to slow (A) and fast (B) brush in peroneal nerve unit. C) Vigorous afterdischarge in CT afferent in response to brush stimulation. The figure shows several consecutive brush strokes. D) Delayed acceleration of impulse response to sustained monofilament indentation (45 mN) in peroneal nerve unit. The initial few seconds of adaptation was followed by, a period of low activity for 20 s, after which firing markedly increased again for about 90 seconds. Note that individual spikes are not discernable and almost completely covers signal background. The inset shows individual spikes superimposed (bar below spikes denotes time 1ms). Arrows denote on and off stimulation where an increased firing is seen as the indentation stops. Panels in (E-G) shows single unit examples of firing across units in the different nerves in response to brush stroking velocity (0.1cm/s - 30 cm/s) and fitted quadratic curve. E) lateral anthebrachial nerve, F) radial nerve, G) peroneal nerve.

After establishing the prevalence of CT units in these nerves and finding that they display similar mechanical response properties in the lower leg as in the arm, our next outstanding question was to explore their sensitivity to chemical agents. Although we had no direct indication from previous studies that CT afferents, or their mouse homologue CLTMs, are sensitive to chemicals, a formal exploration in humans was lacking. We therefore here chose to evaluate the responses to two well-known chemical agents: menthol and histamine.

### Menthol

We first examined the response to menthol application on the receptive field of eight CT units recorded in the arm. For all experiments we initially applied ethanol as a control on the receptive field. We also noted the participants perception of cooling throughout the experiment. There was no activity in CT afferents in response to menthol alone but we could confirm the effect of menthol by recording firing from a background C-temperature sensitive neuron (see arrow denoting smaller unit figure 3C). The participants reported the sensation of cooling and for several recordings a background cooling unit near the recording electrode, whose activity did not correlate with mechanical stimulation, would appear at latency that matched the perception of cooling sensation. We also analyzed whether the firing frequency in response to brushing was altered by the menthol in 5 units. There was no indication that the fast (10 cm/s) and slow (1 cm/s) strokes had any different effect and we therefore pooled the data from these brush velocities. We found no clear indication that menthol modulated the firing rate in response to brush stroking before (Figure 3A) and after menthol application (Fig 3B). The mean of all unit firing in response to brush before menthol application (pre) was 61 Hz, SE +/- 1.9 Hz and after (post) was 50 Hz, SE +/- 2.4 Hz. The individual unit firing in response to brushing pre and post menthol application is visualized in Fig 3D. If anything, there was a trend towards a decrease in firing rate over time in these experiments, which is expected as a result of fatigue to repeated brush stimuli.

**Figure 3.**
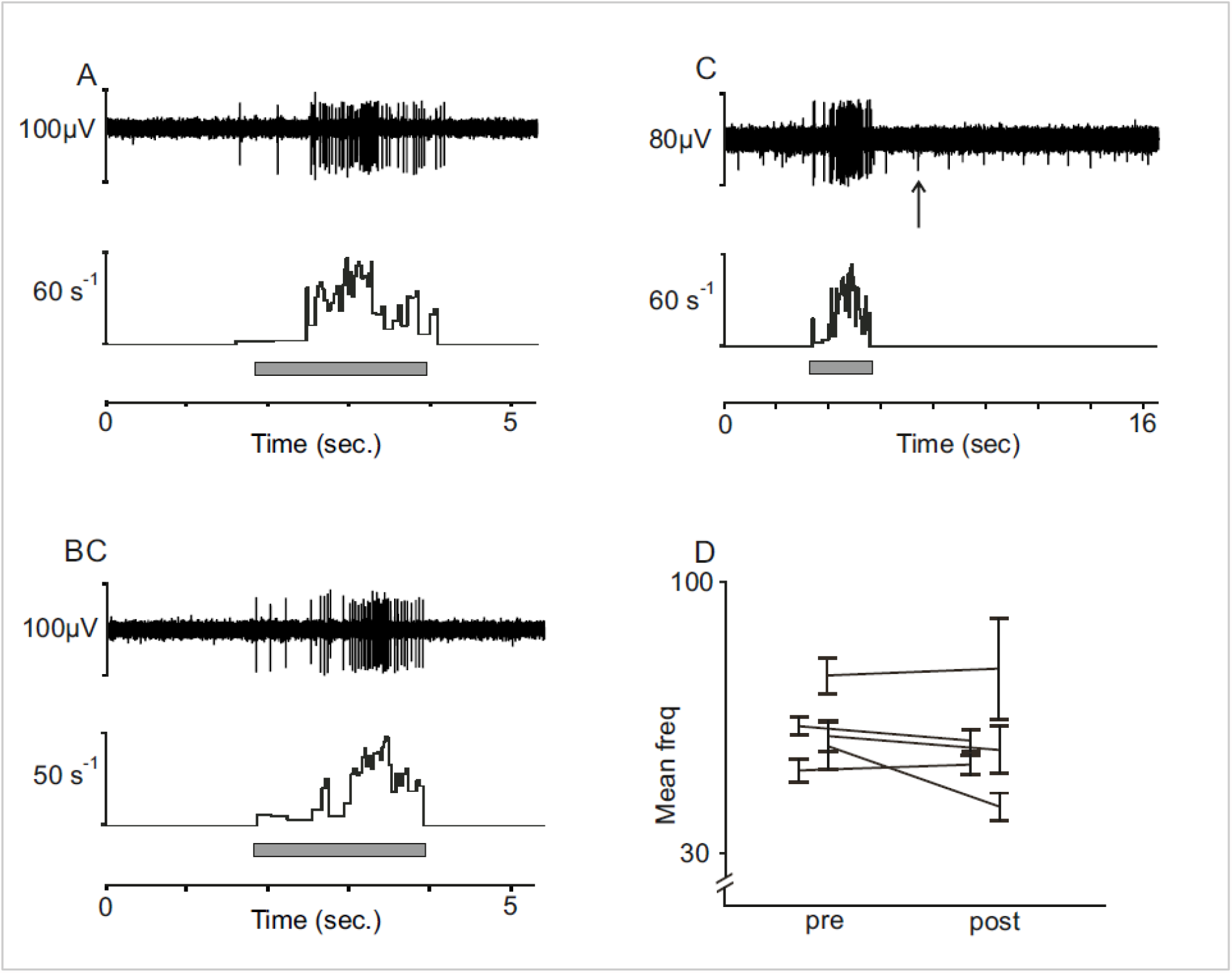
CT afferent response to menthol. A-B) Tactile stimulation with brushing at 1cm/s of CT afferent before (A) and after (C) menthol application. B) Cooling unit in background responding to menthol at time of cooling percept is indicated by arrow. D) Mean and standard error (SE) of firing rate for individual units before (pre) and after (post) application (Mean of all units pre = 61 Hz, SE +/- 1.9 Hz and post 50 Hz, SE +/- 2.4 Hz).

### Histamine

We explored the reaction to histamine iontophoresis in 5 CT units in the nerves of the arm. CT units were characterized as above, and a positive reaction to the histamine was confirmed by a clear flare, and occasionally wheal, on the skin region surrounding the units after iontophoresis. Figure 4B shows a typical flare reaction on the skin surrounding the receptive field of a CT afferent. Iontophoresis of histamine did not evoke any activity in any of the 5 CT afferents. To further assess the effect of histamine on CT afferents, we tested whether the response to a mechanical stimulus was altered by the application of histamine by comparing the response to brush stroking before and after application. We applied fast (10 cm/s) and slow (1cm/s) brush strokes over the course of the experiment. In 2 of these units we were able to keep a stable recording throughout this extended protocol. Figure 4 shows a typical, vigorous response to slow brushing (1cm/s) in one of these units that is similar before (Figure 4A) and after (Figure 4B) histamine iontophoresis. There was no indication that the fast (10 cm/s) and slow (1 cm/s) strokes had any different effect and we therefore pooled the data from these brush velocities. Figure 4C illustrates an example of a flare response to the histamine and Figure 4D illustrates the mean and standard error of firing rate in response to brushing for each unit before and after iontophoresis (mean of pooled units before (pre) = 28 Hz, SE +/- 3.8 Hz and after (post) = 23 Hz, SE +/- 3.3 Hz). We found no indication from these units that mechanical responses were altered by the histamine.

**Figure 4.**
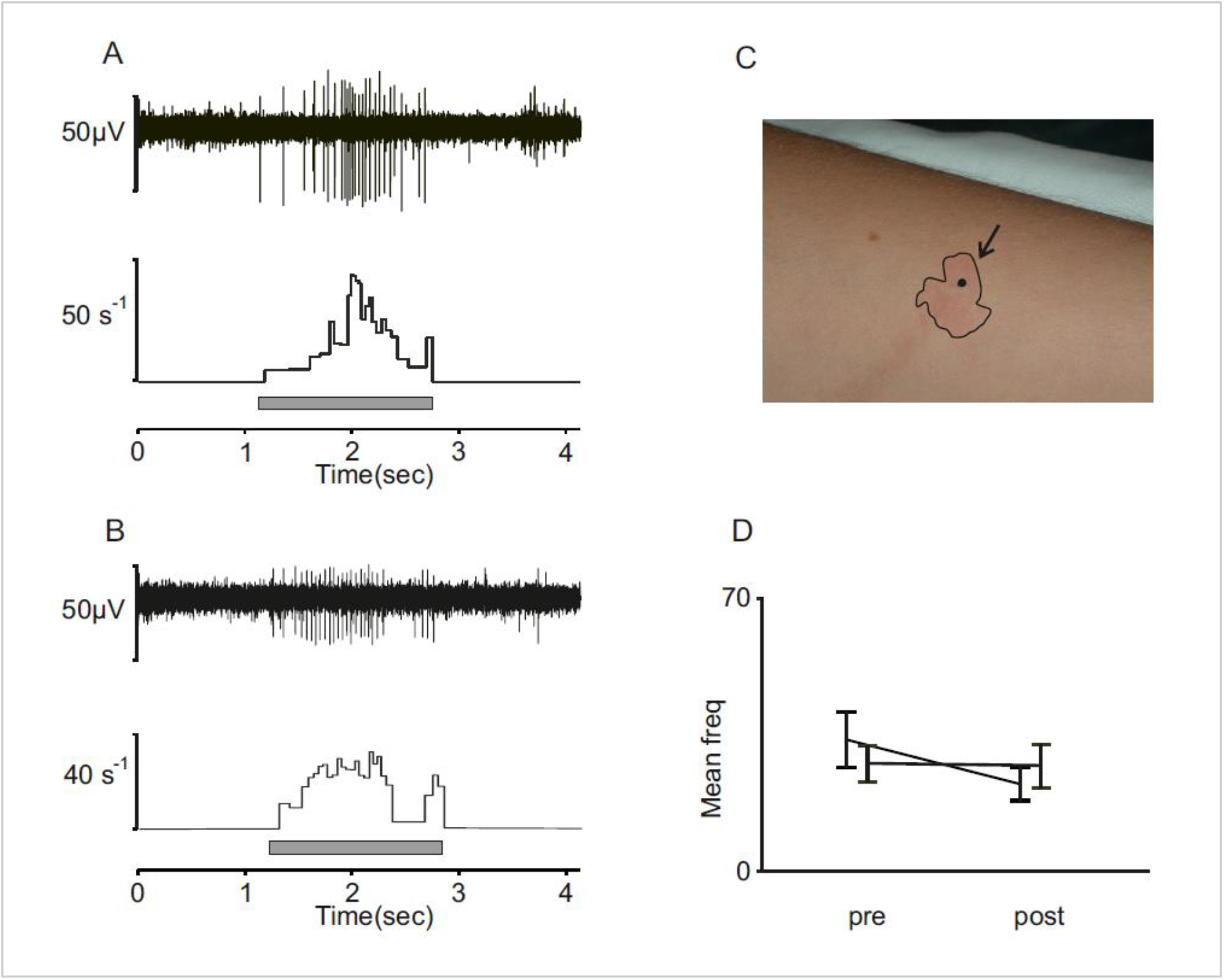
CT afferent response to histamine. A) Response to slow brushing (1 cm/s) in CT unit before iontophoresis. B) Single unit example of response to the same stimulus after histamine iontophoresis. C) Wheal and flare to confirm reaction. D) Mean and standard error (SE) of firing rate for individual units before (pre) and after (post) iontophoresis (Mean of both units pre = 28 Hz, SE +/- 3.8 Hz and post 23 Hz, SE +/- 3.3 Hz).

We also documented the participant’s perception starting at 2 minutes post histamine iontophoresis and continuing at 3-minute intervals for 20 minutes following the histamine iontophoresis. Although the concentration used evoked a clear reaction in the skin, and typically evokes itch sensation (Schmelz et al. 1997), the participants did not here report any clear sensation in relation to the histamine 3 minutes after application. The size of the flares were measured and ranged from 15-22 mm at their maximum. In the example shown in Figure 4B the flare had a diameter of 13 mm at 3 minutes after iontophoresis and extended to maximum 15 mm.

## DISCUSSION

Despite the overwhelming evidence that CT afferents are present across the hairy skin of mammals, a comprehensive account of these afferents presence across different skin sites in man has been lacking. For the first time, we here describe the presence of CT afferents in the peroneal nerve and compare their incidence in microneurographic recordings to that of the lateral antebrachial and radial nerves. The most distal receptive fields were located on the proximal phalanx of the third finger for the superficial branch of the radial nerve, and approximately six centimeters proximal to the lateral malleolus for the peroneal nerve. CT units also responded in a similar fashion across the antebrachial, radial and peroneal nerves with regard to their physiological properties such as conduction velocity, mechanical threshold and in their tuning to brush velocity.

Despite numerous microneurographic recordings from the peroneal nerve, CTs have not previously been reported in this nerve. The reason is probably in part because most microneurography studies of the peroneal nerve have relied on distinguishing C-mechanoreceptive afferents based on intrinsic properties of axonal conduction latency changes in response to repetitive electrical stimulation or combined with natural stimuli, a method known as the “marking technique” (Schmelz et al. 1995; Schmidt et al. 1995; Serra et al. 1999). Originally well validated and utilized for C mechano-nociceptors (CMs), it was only recently adapted to detect non-nociceptive CTs (Watkins et al. 2017). Here we did an extensive experimental series where we actively searched for and identified four CT afferents in the lower leg. Aside from their low-threshold and slow conduction velocity, their vigorous response to brushing and their tuning to brush velocity was consistent with the response properties of CT units described in the arm. One of the peroneal CT units also exhibited delayed acceleration to sustained indentation of a von Frey filament. The phenomenon has been described for C-tactile afferents in the arm (Vallbo et al. 1999) and, in this as well as in previous recordings, is characterized by an initial few seconds of adaptation followed by a period of low activity after which firing markedly increases again for ∼20 seconds. An off-response characterized by a slight increase in firing as the indentation stops is typical and was also seen in this unit. Considering the number of experiments (17) where we actively searched for CT units in the peroneal nerve, the number of CT units identified was relatively low compared to that in experiments for the arm. An important factor that influences the incidence of identifying different afferents in single unit recordings is the search technique. When the same technique searching for low threshold mechanoreceptors in the nerves of the arm, such as a light stroking of the skin, identifying CT afferents is much more common and we are as likely to identify CT afferents as we are to record slowly adapting type 1 units (SA1). This prevalence of CT afferents relative to SA1 has been consistent across different studies (Ackerley et al. 2018; Loken et al. 2009; Vallbo et al. 1999; Wessberg et al. 2003). Part of the difficulty identifying CT afferents in experiments of the peroneal nerve was that sympathetic background activity (Macefield and Wallin 2018) was more common than in the skin nerve branches of the arm. Although readily identified as efferent activity, we decided to actively compare the incidence of CT afferents with high threshold C-fibres responding to pinching the skin. However, when applying a natural stimulation search technique, we found that CT afferents may indeed be sparser than CMs in the lower leg. When actively stroking or pinching the skin to detect CT and CM respectively, we recorded from CMs 5 times more often than CTs (see figure 1). These estimates are obviously not an exact measure of density, but as our main focus was to record from CT afferents, the higher incidence of CMs in relation to CTs is more likely to be an underestimation, than vice versa. We recently showed that low-threshold mechano C afferents are also found in glabrous skin of the human hand, although their presence here appear to be very sparse (Watkins et al. 2021). It is now clear that CT afferents can indeed be found in the distal parts of the limbs such as the hand and lower leg, although the current results suggests that they are considerably sparser in the distal parts of the human body, similar to what has been found in non-human primates (Kumazawa and Perl 1977)

CT afferents display several interesting physiological phenomena that are yet to be fully understood. We here resumed investigation of some of the early observations made on these afferents. A common feature is afterdischarge, characterized by a vigorous response to a natural stimulus where a trail of impulses follow a natural stimulation. This phenomenon is more prevalent when the stimulus is novel and therefore often appears just as the unit is being identified. It can be difficult to reproduce during recordings after the unit is subjected to more stimuli. Afterdischarge is not unique to CT afferents but is also observed in C-nociceptors (Iggo 1960). In our sample, 4 out of 20 nociceptor units displayed this feature upon mechanical threshold identification. In our previous observations afterdischarge in CT afferents is typically halted by re-applying a stimulus, but we here show that this is not always the case. Our example (Figure 2C) shows a CT unit that continuously afterdischarges in response to brushing repeated at slow and intermediate velocity. Whether this continuous afterdischarge was a unique feature of a single afferent or whether it can be triggered during particular circumstances is not known but warrants further study. A more unusual feature is delayed acceleration. Previous studies have reported that delayed acceleration is occasionally observed in CT units of the lateral antebrachial cutaneous nerve (Vallbo et al. 1999) and can be reproduced in the units that display this feature. Besides the peroneal nerve unit that displayed delayed acceleration in our example, we found this phenomenon in two units tested in the radial and antebrachial nerves. Neither afterdischarge nor delayed acceleration appear to have any clear perceptual correlate and the functional significance of this type of response is unknown. However, any perceptual correlate is also likely dependent on spatial summation of units displaying these features simultaneously which has so far been precluded in single unit recording studies alone.

We chose to continue our explorations to include the cooling agent menthol and histamine, a well-known pruritogen. Classification of sensory neurons by RNA sequencing show that C low-threshold mechanoreceptors in rodents are clearly distinguishable from nociceptors and that they do not express the TRPV1 receptor (Usoskin et al. 2015), a finding that has been confirmed by the lack of response to heat in human microneurography recordings (Nordin 1990; Vallbo et al. 1999). Animal studies have also suggested that CLTMs respond to rapid cooling (Bessou et al. 1971; Hensel et al. 1960; Seal et al. 2009). Nordin (Nordin 1990) found that some CTs responded with a short burst of spikes to cooling, but most of these studies have included a light mechanical impact on the receptive field with the result that the response to pure cooling has been somewhat elusive. Despite their insensitivity to heat, previous research suggests that the responses of CT afferents to mechanical stimuli can be modified by temperature. For example, findings from microneurography suggests that the responses of CT afferents are optimal at skin temperature (Ackerley et al. 2014). Furthermore, they are modified by temperature such that touch above skin temperature decreases their firing and cool touch lower their firing but often produces a longer lasting firing at low frequency, i.e. afterdischarges (Ackerley et al. 2018). However, the mechanisms underlying the responses of CT afferents in combination with cooling are still unknown. To add to our understanding, it was therefore of great interest to find out whether the cooling agent menthol, that is dependent on the thermosensitive cation channel TRPM8, in itself or in combination with brushing, acts on C-tactile afferents. In the 8 units where menthol was applied, none were directly activated by menthol. In a subset of units we also did repeated tests of brushing at different velocities before and after menthol application, but found no indication that the firing rate was modulated by menthol at the time of cooling percept. In fact, there was no indication of a decrease in firing with repeated tests beyond what can be attributed to normal variability and fatigue to repeated stimuli. Our finding that menthol does not directly act on CT afferents is consistent with data from rodents suggesting that TRPM8 is not expressed in CLTMs but instead mark a highly selective class of primary afferent neurons (Usoskin et al. 2015).

Although there are various substances that induce itch, histamine is one of the best-known endogenous substances. Histamine acts directly on a prurigenic class of primary sensory neurons containing calcitonin gene-related peptide- and substance P (Schmelz et al. 1997; Tani et al. 1990). Histamine induced itch is primarily mediated by mechano-insensitive C-fibers (MIA) that are also sensitive to heat or capsaicin. Studies from mice show that sensory neurons that respond to histamine include those that express Mas-related G-protein coupled receptor member A3 (MrgprA3). However, histamine can also signal through TRPV1 and in primates activates mechanosensitive sensory neurons such as A-fiber mechano-heat (AMH) and C-fiber mechano-heat (CMH), although to a weaker extent than MIA (LaMotte et al. 2014). None of the CT units in our sample could be directly activated by histamine iontophoresis. Our results are consistent with one of the earliest reports on the physiological properties of low threshold C-afferents in cat where histamine did not provoke a consistent response even at concentrations (30% in saline scratched into the skin) that would cause wheal in both human and cat skin and also cause scratching in man (Bessou et al. 1971). Because cooling has been reported to modulate the sensitivity of CT afferents to mechanical stimuli we also tested if histamine had a similar effect on response to brushing in a subset of these units. Brushing the receptive field after histamine iontophoresis had no clear effect on brushing sensitivity, and if anything, the responses to brushing decreased. Histamine is only one of many pruritogens, and in spite of its popularity in experimental studies, antihistamines are seldom effective as relief in clinical itch populations. The accumulated knowledge from our studies on humans and other species strongly suggest that CT afferents lack both the receptor for histamine and TRPV1 and that they do not play a role in histaminergic itch. However, some inconsistencies remain regarding their chemical responsiveness. For example, early recordings in CLTMs in cat showed no response to application of acetic acid (Bessou et al. 1971), which acts on the TRPA1 receptor (Wang et al. 2011), while recent single cell sequencing from rodents suggest that TRPA1 may be expressed to some extent in CLTMs (Usoskin et al. 2015). The possibility also remains that CT afferents play a role in touch evoked itch, alloknesis, which is yet to be addressed in detail.

Molecular visualization of the apparent CT afferent mouse homologue have shown that CT afferents are present in all the nerves supplying the hairy skin (Li et al. 2011). Combined with ours and previous recordings from nerves innervating the skin of the face, thigh and palm (Edin 2001; Johansson et al. 1988; Nordin 1990; Watkins et al. 2021), this suggests that CT afferents are ubiquitous across human hairy skin although very sparse in distal parts of the body. Deep sequencing of sensory neurons from mice have revealed that while the most dramatic differences in gene expression patterns are observed between unmyelinated and myelinated neuron subtypes, some genes such as the potassium voltage-gated channel Kv4.3 is uniquely expressed in CLTMR sensory neurons (Zheng et al. 2019). Although the possibility that there are important differences between the species should be considered, the data from mouse and humans all point in the direction that CT afferents in man and C low-threshold afferents in mice to a large extent exhibit similar physiological properties. In light of previous studies on the tuning to brush velocity (Loken et al. 2009), the combined effects of temperature and mechanical stimulation (Ackerley et al. 2014; Ackerley et al. 2018) and the lack of activation by chemical agents alone (Bessou et al. 1971), our results adds to the growing body of research from several species suggesting that CT afferents constitute a unique population of cutaneous mechanosensory afferents with highly specialized mechanisms optimal for transducing features of the gentle caressing stimulation seen in affiliative behaviours.

Detection of touch requires specialized receptors with unique morphological, chemical and mechanical properties that allow us to sense the different modalities of touch. A long-debated question has been to determine if a particular type of afferent neuron is responsible for a given sensation, or if sensation is the composite result of contributions from a diverse population of primary afferents (Braz et al. 2014; Craig 2003; Ma 2010; Perl 2007). Present evidence suggests that primary afferents are highly specialized to detect different aspects of external stimuli. However, research from both humans and rodents suggest that integration between the coding properties of several types of afferents is necessary for the percept of for example mechanical allodynia (Löken et al. 2017) and temperature (Paricio-Montesinos et al. 2020). The precise contribution of other fiber types for the coding of affiliative hedonic touch is not known, but clearly also depends on Aβ afferents for conscious sensation (Olausson et al. 2002), as well as being heavily modulated by factors such as context and homeostatic state (Kringelbach and Berridge 2017).

## Acknowledgements

We thank Karin Göthner for excellent technical assistance and the participants for their patience.

## Funding

Åke Wiberg Foundation to Line Löken and Swedish Research Council 2017-01717; Västra Götaland Region ALFGBG-725751 to Johan Wessberg.

## Supplementary figure legends

**Supplementary figure 1**.

Example of afterdischarge in a C unit with high mechanical threshold (3g) in response to a sharp mechanical probe. Lower panel indicates timing of probe stimulation and the firing continues for several seconds after the probe is removed.

## Notes

### Competing Interest Statement

The authors have declared no competing interest.

