## Supplementary figures and images for "A topographical and physiological exploration of C-tactile afferents and their response to menthol and histamine"

### Supplementary figure 1

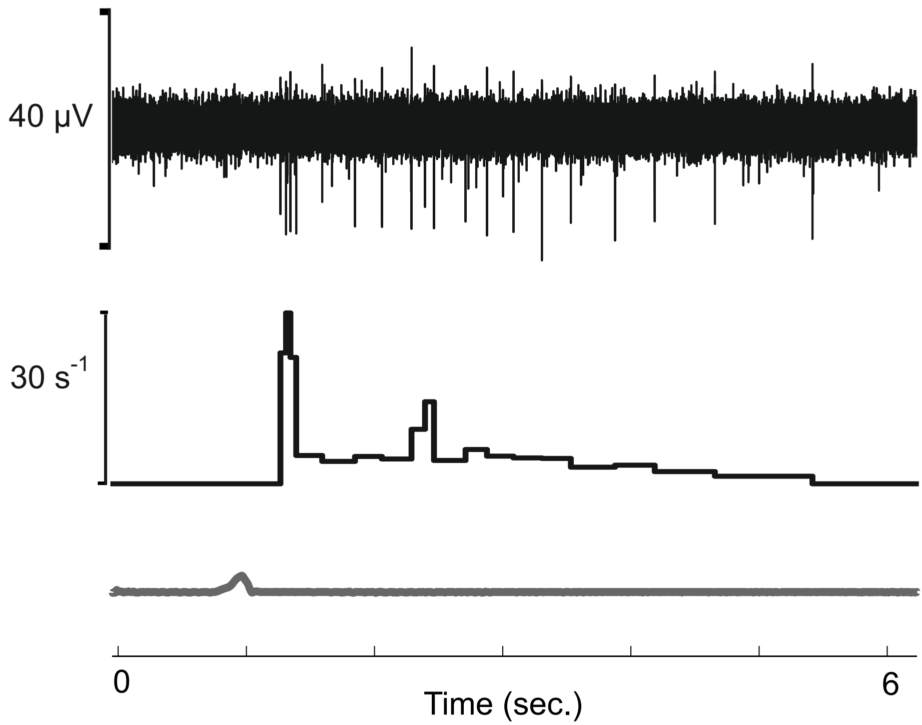
